# Mapping interactions between disordered regions reveals promiscuity in biomolecular condensate formation

**DOI:** 10.1101/2023.07.04.547715

**Authors:** Atar Gilat, Benjamin Dubrueil, Emmanuel D. Levy

**Affiliations:** Department of Chemical and Structural Biology, Weizmann Institute of Science, Rehovot, Israel

## Abstract

Intrinsically-disordered regions (IDRs) promote intracellular phase separation and the formation of biomolecular condensates through interactions encoded in their primary sequence. While these condensates form spatially distinct assemblies in cells, it is unclear whether such specificity can be conferred by IDRs alone. Indeed, IDRs exhibit high conformational flexibility whereas specificity in protein recognition is generally associated with well-defined 3D structures. To characterize IDR-IDR interactions and assess their ability to mediate self-specific partitioning, we developed a synthetic system of Multivalent IDRs forming Constitutive DROPlets (*micDROP*). We investigated ten natural IDRs that underwent phase separation in *micDROP*. These IDRs exhibited a wide range of saturation concentrations *in vivo*, which correlated well with their total sequence stickiness. We then probed IDR-IDR specificity by co-expressing pairs of IDRs fused to homologous scaffolds that did not co-assemble. We observed a high degree of promiscuity, particularly among IDRs from the proteins Ddx4, DYRK3, ERα, FUS, hnRNPA1, HspB8, RBM14 and TAF15, whereas TDP43 and UBQ2 formed spatially distinct condensates regardless of their partner. Further investigation revealed the short and conserved α-helical segment of TDP43’s IDR was governing its specific self-recognition. Our findings imply that IDRs can tune their phase separation propensity through sequence composition, while their formation of discrete condensates likely requires additional cellular or structural determinants.

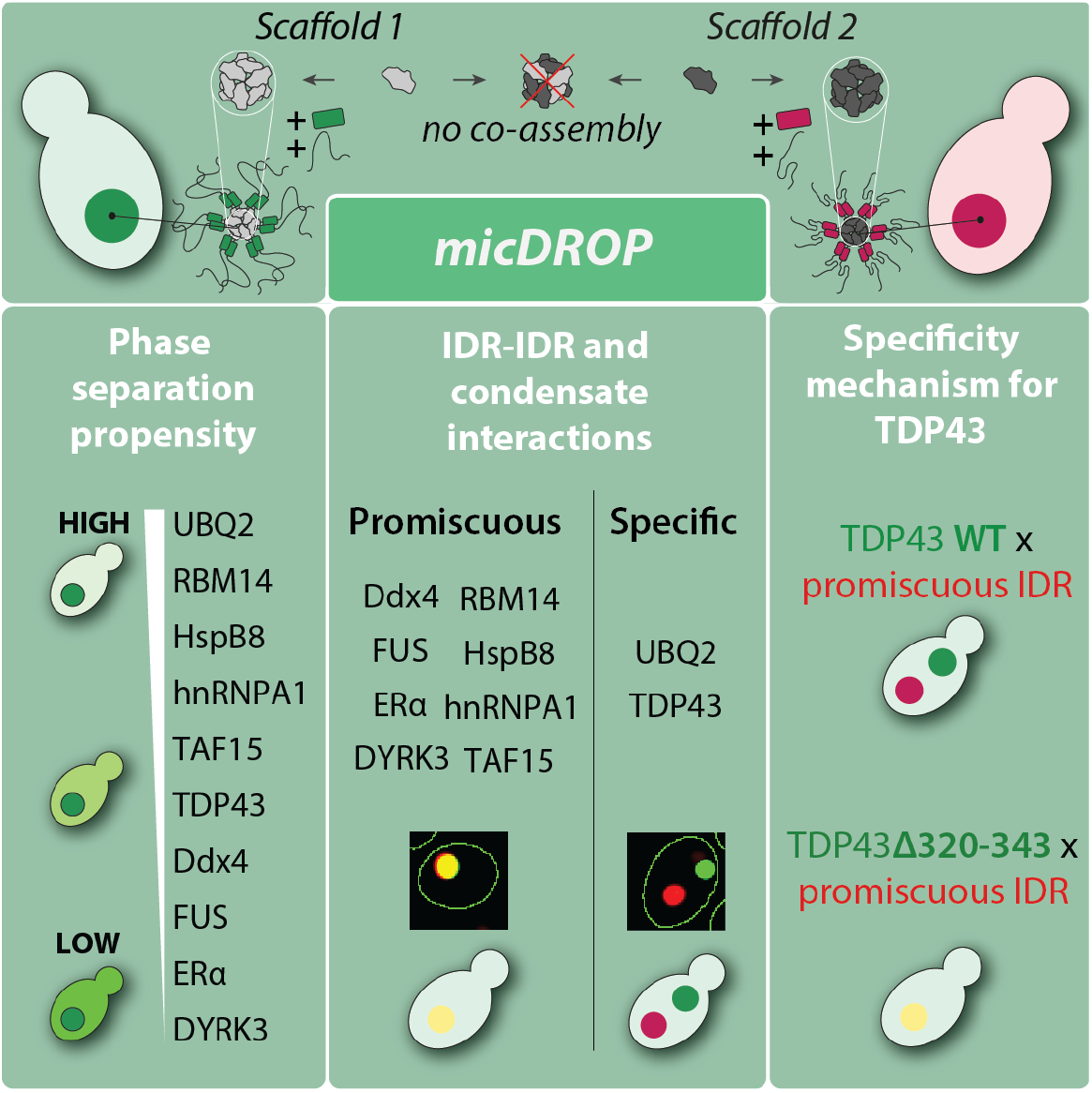

## INTRODUCTION

Intrinsically disordered regions (IDRs) represent a unique class of protein segments that lack a well-defined tertiary structure under physiological conditions. These regions possess inherent flexibility, enabling them to sample a variety of conformations and engage in diverse interactions^1^. IDRs are prevalent in eukaryotic proteomes and play crucial roles in a wide range of biological processes^2^. One of the key emerging functions of IDRs is in mediating the formation of biomolecular condensates. Condensates are membraneless compartments within cells that facilitate the spatial organization of biomolecules and regulate a variety of cellular processes related to stress responses^3^, cellular mitosis^4^, DNA replication and repair^5^, and more^6–9^. Condensates may form through phase separation, whereby IDRs self-interact and demix from the surrounding bulk (cytoplasm/nucleoplasm), and the resulting assemblies often display liquid-like behavior^9–11^. Experimental works conducted over the past decade have consistently shown that, for many protein components of biomolecular condensates, the IDR sequence alone can undergo phase separation through homotypic interactions both *in vitro* and *in vivo*^*12–14*^.

Each type of condensate has a unique milieu of components^15–17^. However, proteins such as FUS (FUsed in Sarcoma) have been identified in functionally and spatially distinct condensates including stress granules^18^, nucleoli^19^ and paraspeckles^20^. The ability to partition proteins into discrete condensates without their intermixing implies some measure of specificity in the interactions driving their formation. Whether such specificity can be conferred through the IDR sequence alone, or whether additional cellular factors are required, remains unclear.

In the absence of a stable structure, the amino acid composition of an IDR sequence determines its ability to phase-separate, particularly through aromatic or charge-driven interactions^21–28^. Unlike protein folding, in which the sequence itself is critical, overall compositional characteristics can explain an IDR’s phase separation behavior. For example, the overall number of aromatic residues and their spacing modulate hnRNPA1’s propensity to phase-separate^29^. This notion is in line with the fact that only ∼5% of amino acid positions in IDR sequences are conserved across orthologs^30^, while other molecular features, such as sequence length or net charge, are under evolutionary constraint. At the same time, it is unclear whether amino acid composition alone can confer specificity in IDR-IDR interactions because interaction energies between amino acids show that no pair is exclusively favorable^31^.

To further our understanding of possible IDR-IDR specificity in intracellular phase separation, we present *micDROP* (Multivalent IDRs forming Constitutive DROPlets), a synthetic system wherein two non-interacting dodecameric scaffolds present different IDRs and are co-expressed in yeast cells. Using confocal fluorescence microscopy, we quantitatively assessed the propensity of nineteen candidate IDRs to undergo phase separation *in vivo*. Our findings revealed that IDR sequences underwent phase separation with a wide range of apparent affinities, as inferred from their relative saturation concentration (C_sat_). Despite displaying a wide range of C_sat_ and sequence characteristics, most IDRs interacted with high promiscuity and colocalized indiscriminately. Two IDRs, UBQ2 and TDP43, formed self-specific condensates. Examining TDP43’s IDR revealed that a transient α-helical segment governed this self-specificity. Indeed, deletion of this short segment did not impact TDP43’s ability to phase separate, but abolished its specific self-recognition. Overall, *micDROP* provides a novel method for the quantitative study of phase separation *in vivo* and may elucidate additional features important both for phase separation propensity and specificity.

## RESULTS

### micDROP: A synthetic system for studying IDR-mediated phase separation in vivo

The genetic construct of *micDROP* consists of three domains connected by flexible linkers: a scaffold subunit, a fluorescent reporter, and an IDR of interest (**Figure 1A**). This design bears similarities to *Corelets*, an optogenetics-based system previously introduced by Bracha et al.^32^, with two significant distinctions: *micDROP* relies on constitutive oligomerization, and utilizes a different scaffold. Indeed, in our initial experiments, the 24-mer ‘maxi’ ferritins tested *in vivo* were prone to aggregation (**Figure S1**), prompting us to select 12-mer ‘mini’ ferritins^33^, hereafter referred to as Dps (DNA Protection during Starvation) scaffolds. The resulting fusion protein (**Figure 1B**) forms a dodecameric complex (**Figure 1C**), thus increasing the effective valency of the IDR. This fusion strategy was selected because many IDRs, such as NPM1^34^, FUS^35^, and TDP43^36^ exist in homo-oligomeric proteins. Moreover, a higher valence is known to enhance phase separation^37–39^, which we expected would enable tracking the phase separation behavior of IDRs with weak propensities for homotypic interactions. *micDROP*-green, consisting of a Dps scaffold from *V. cholerae* (PDB code 3IQ1) fused to a yellow fluorescent protein (Venus^40^), was first tested with the disordered region of hnRNPA1, an RNA-binding protein (RBP) whose phase separation propensity is well-documented^22,29^. Both the monomeric hnRNPA1 IDR as well as the IDR-free *micDROP*-green were homogeneously dispersed when expressed in yeast cells. However, hnRNPA1 fused to *micDROP*-green formed bright puncta, indicating the formation of synthetic condensates (**Figure 1D**).

**Figure 1.**
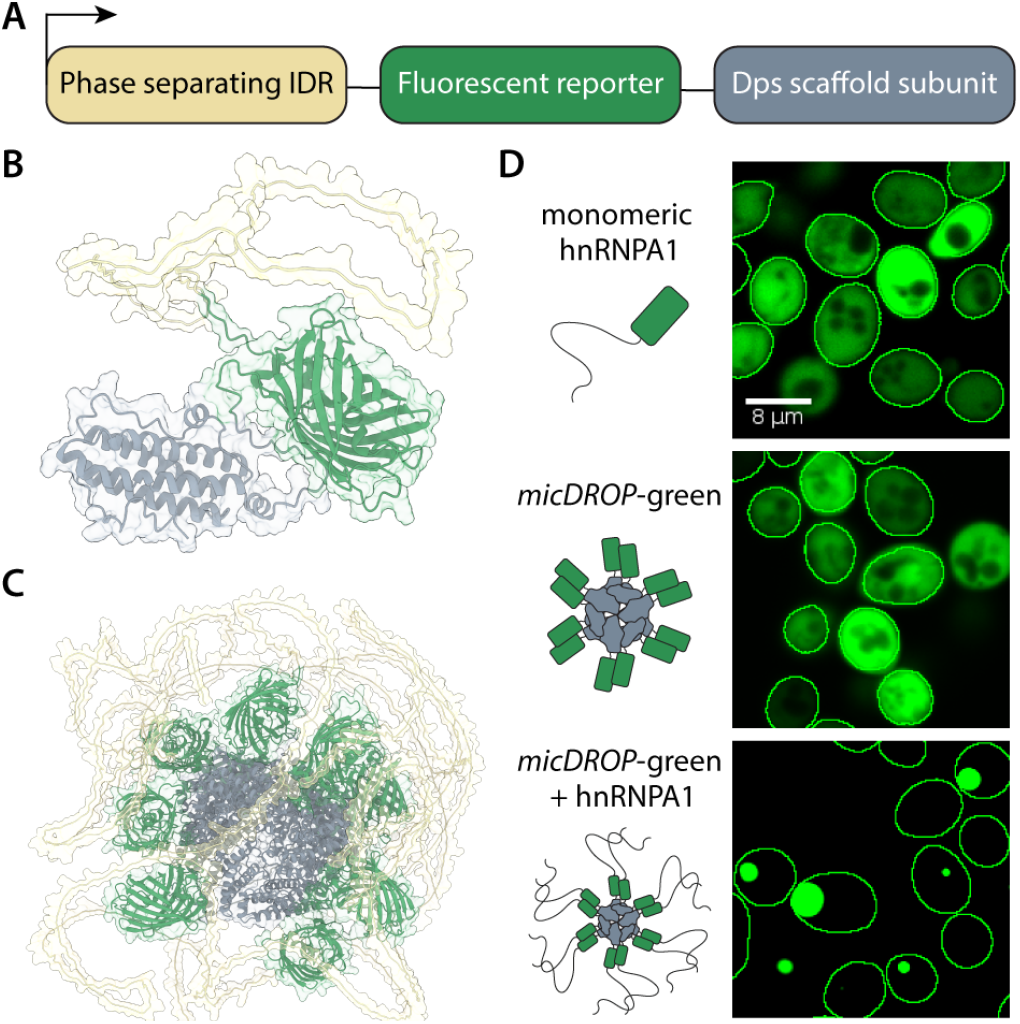
*micDROP*: A synthetic system for studying IDR-mediated phase separation *in vivo*. **A**. Schematic representation of *micDROP*’s genetic construct. It comprises three domains connected by flexible linkers: an IDR of interest (yellow), a fluorescent reporter (green), and a Dps subunit as a homo-oligomeric scaffold (gray), which assembles into a dodecamer. **B**. Cartoon rendering of a single fusion protein harboring an IDR region. The structure model was generated with AlphaFold2^41^. **C**. Cartoon rendering of an assembled complex model generated with AlphaFold2. Two fluorescent proteins obstructing the view of the scaffold, as well as two subunits in the back of the structure, were hidden for clarity. **D**. Fluorescence microscopy images of yeast cells expressing three constructs: a monomeric construct encoding an IDR region from hnRNPA1 fused to Venus (top), *micDROP*-green - an assembled Dps complex from *V. cholerae* fused to Venus without an IDR region (center), and *micDROP*-green fused to the IDR region from hnRNPA1 (bottom).

### micDROP captures IDR propensity for phase separation

Phase separation is characterized by a phase boundary, which reflects the set of conditions at which demixing occurs (**Figure 2A**). In living cells, where temperature and pressure remain relatively constant, phase separation can be modulated by intracellular conditions such as crowding or pH, or component-related parameters, including multivalency, affinity, and concentration^42^. Because *micDROP* has a fixed valency and is expressed in similar conditions, the primary parameters modulating the phase boundary ought to be the intracellular concentration of the system and the homotypic affinity of the IDR. We sought to identify whether a phase boundary existed - i.e., a saturation concentration (C_sat_) beyond which *micDROP* would phase separate into a dense phase, visible as a bright puncta, and a dilute phase, observable as homogeneous fluorescence in the cytosol. If a C_sat_ existed for an IDR in *micDROP*, we hypothesized its value should reflect the apparent affinity of homotypic interactions for that IDR.

**Figure 2.**
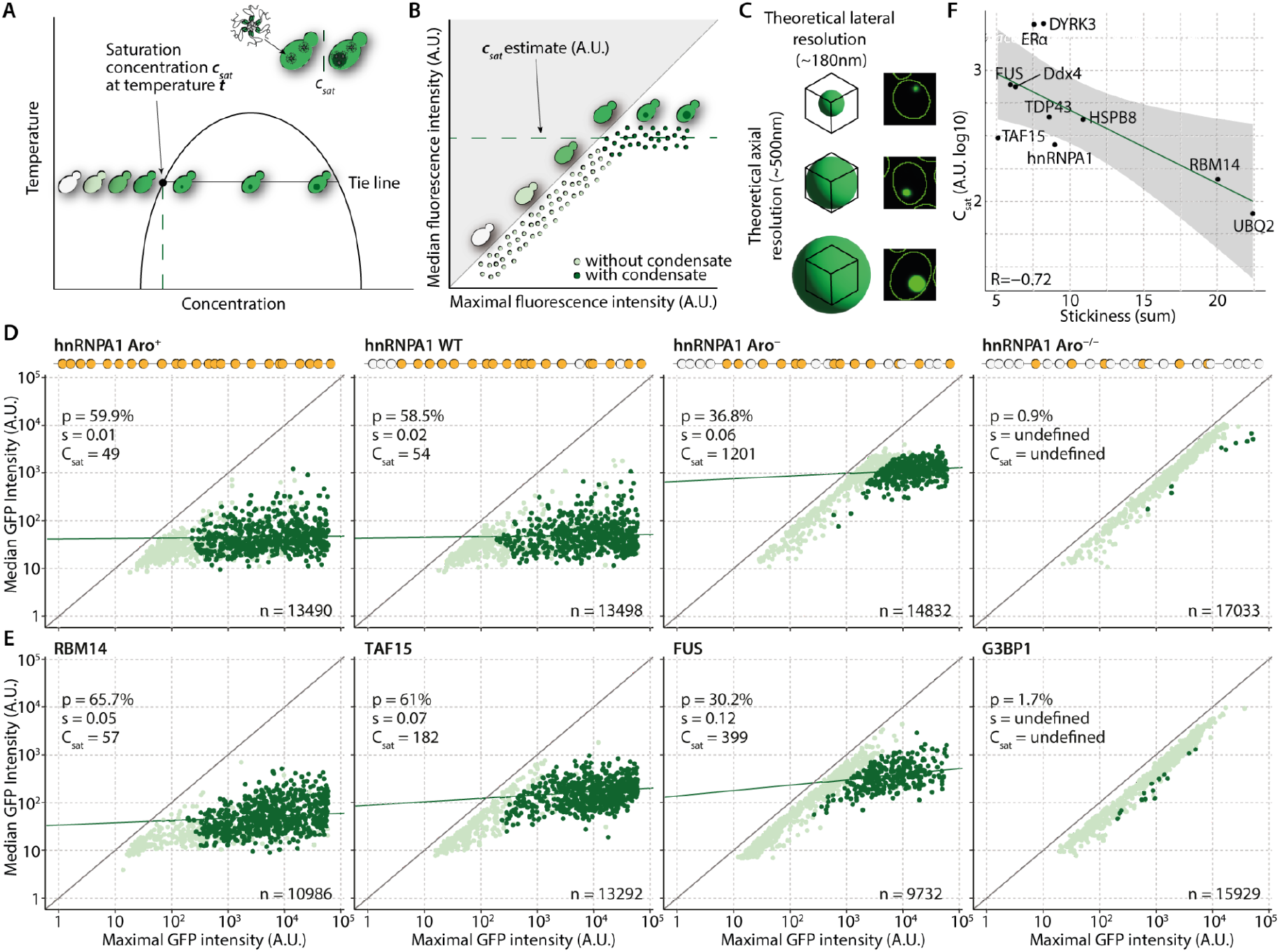
*In vivo* quantification of C_sat_ for model IDRs. **A**. Schematic representation of a phase diagram. At constant temperature (*t*), increasing the concentration to the point of saturation (C_sat_) will result in phase separation into a condensate (visible as a bright puncta) and a dilute phase. Further increase of concentration will increase condensate volume without increasing its concentration. **B**. Schematic representation of a method to estimate C_sat_ from the fluorescence intensity of the dilute phase (cytoplasm) among cells with a detectable dense phase (condensate or puncta). Each segmented cell’s median fluorescence intensity is recorded (y-axis) and reflects the dilute phase concentration. This quantity is analyzed as a function of the maximal fluorescence intensity in the same cell. Cells without a condensate (light green) show maximal intensities equal to ∼1.5-fold their cytosolic median intensity. By contrast, cells with condensates (dark green) show maximal fluorescence intensities 3- to 100-fold larger than the median. The dilute phase concentration among these condensate-containing cells reflects the C_sat_. **C**. The size of condensates varies while the confocal volume is fixed, making it difficult to measure protein concentration in the dense phase: small condensates do not fill up the entire volume and their fluorescence intensity is underestimated, while condensates larger than the confocal volume will appear brighter due to fluorescence leakage. Consequently, cells with condensates display a wide range of maximal intensities associated with condensate size, but relatively constant median intensities (reflecting C_sat_). **D**. Modulation of hnRNPA1 C_sat_ through changes in aromatic residue content. Three hnRNPA1 mutants previously characterized *in vitro*^*29*^ were expressed with *micDROP*-green and their relative C_sat_ measured. Aromatic residues are represented by orange dots in the cartoon sequences above each scatterplot. The fraction of cells identified as containing a condensate is measured (*p*), along with C_sat_ and the slope (*s*) of the linear regression. To facilitate comparison we show the same number of randomly sampled data points (n=1000) for each scatterplot. Decreased aromatic residue content inhibits phase separation and increases C_sat_. **E**. C_sat_ measurements of three model IDRs (RBM14, TAF15, and FUS) using *micDROP*-green. G3BP1’s IDR serves as a negative control and does not form condensates even at high expression levels, in agreement with *in vitro* data^47^. **F**. The total stickiness^31^ of IDRs correlates with the C_sat_ measured *in vivo*, with stickier IDRs exhibiting higher propensities to phase separate.

The first criterion to determine IDR-mediated phase separation in *micDROP* was the presence of condensates in cells, which were identified and segmented using custom scripts for image analysis^43^ (**Methods**). By plotting the median fluorescence intensity against the maximal intensity for each segmented cell, we predicted that cells without a condensate would spread along an approximately linear diagonal and display a constant ratio between median and maximal fluorescence intensity (**Figure 2B**). By contrast, cells with condensates were expected to display constant median and maximal fluorescence intensity values. Indeed, in accordance with phase separation theory, the dense and dilute phases should exhibit constant concentrations; an increase in expression of *micDROP* beyond the phase boundary should therefore increase the dense phase volume, but not its concentration. However, due to limitations of fluorescence microscopy, particularly in resolution^44^, an accurate assessment of the fluorescence intensity within the dense phase was not possible, resulting in a wide range of maximal intensities that reflect condensate size (**Figure 2C**). As measuring the dilute phase (the cytoplasm) was not similarly limited by resolution, we reasoned that the measured median intensity at which condensates began appearing reflected C_sat_. We expected that such inferred C_sat_ should be relatively constant, which we assessed from the slope of a linear regression of C_sat_ values among cells containing condensates. We classified an IDR as undergoing phase separation in *micDROP* when at least 20% of the cells were identified as containing a condensate (p≥20%), and when the slope (s) of the linear regression fitted for those cells was ≤0.4, indicating that beyond the phase boundary, the concentration of the dilute phase remained relatively constant.

To evaluate our ability to determine relative affinity from C_sat_ measurements, we examined how mutations within an IDR sequence affected its propensity to phase-separate. Previous work showed that radii of gyration of monomeric hnRNPA1 IDR and its critical temperature for phase separation could be modulated by the addition or removal of interaction ‘stickers’ in the form of aromatic residues (Tyr/Phe)^29^. We used *micDROP*-green to express three hnRNPA1 mutants previously tested *in vitro*: Aro^+^, with five Ser→Phe mutations, Aro^**–**^, with six Phe→Ser mutations, and Aro^−/−^, with thirteen Phe→Ser mutations (**Figure 2D**). hnRNPA1 Aro^+^ and hnRNPA1 WT showed a comparable propensity to undergo phase separation, with a similar percentage of cells identified with condensates and C_sat_ values. While the calculated C_sat_ for Aro^+^ was slightly lower, the difference did not match previous *in vitro* results, in which a ∼10-fold decrease in critical concentration was observed at 25°C. However, as C_sat_ values were close to autofluorescence levels (∼20 A.U.), it is probable that the measurement of the Aro^+^ mutant C_sat_ was overestimated. By contrast, hnRNPA1 Aro^**–**^ showed a ∼1.6-fold decrease in the percentage of cells containing condensates and a ∼20-fold increase in C_sat_ compared to hnRNPA1 WT, while hnRNPA1 Aro^−/−^ did not undergo phase separation and no condensates were observed even at high expression levels (p<1%). These *in vivo* experiments recapitulated observations previously made *in vitro*, where Aro^**–**^ displayed a ∼15-fold increase in C_sat_ at 25°C, and Aro^−/−^ was not observed to undergo phase separation at any concentration at 25°C. From these experiments, we concluded that *micDROP* captures differences in apparent homotypic affinities of IDR interactions that drive phase separation.

### IDRs display a wide range of phase separation propensities that correlate with their overall stickiness

The validation of *micDROP* on hnRNPA1 mutants motivated us to carry out a broader analysis across various IDR families. We selected eighteen IDR candidates (**Table 1**) referenced in the PhaSePro database^45^ and previously shown to undergo phase separation through homotypic interactions either *in vivo* or *in vitro*. In addition, we included as a negative control the disordered region of G3BP1, a scaffold protein in the assembly of stress granules^46^ that binds RNA. While the C-terminal IDR of G3BP1 was found to be essential for phase separation of the full-length protein, it was unable to drive it without the nearby RNA-recognition motif (RRM) binding to RNA^47^. As such, the G3BP1 IDR was not expected to form condensates in *micDROP*.

**Table 1.**
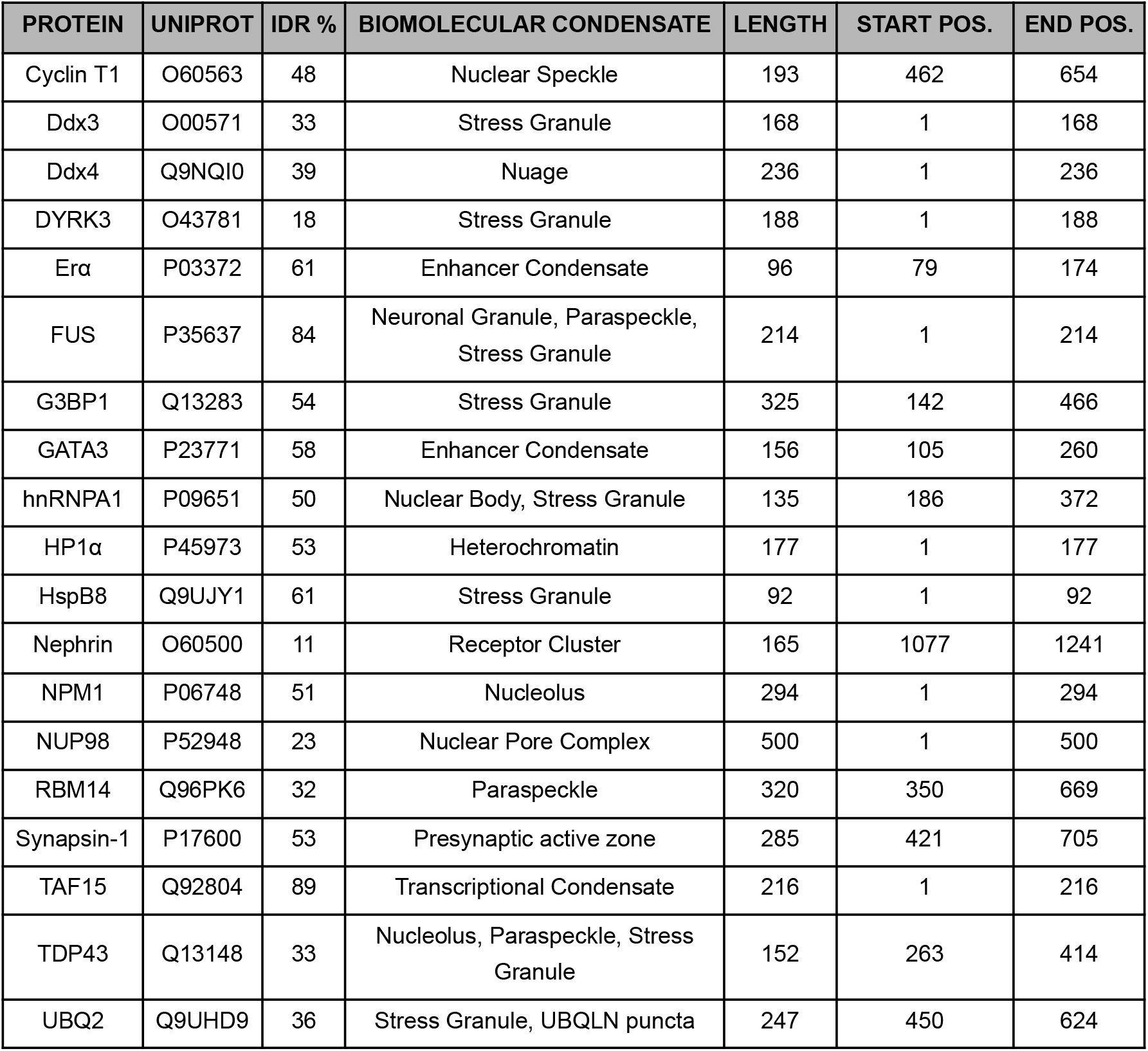
Candidate IDRs for phase separation using *micDROP*. Disordered fraction of the protein was derived from MobiDB^49^ consensus prediction. Biomolecular condensate, start and end positions in sequence were derived from annotations in the PhaSePro database^45^.

Importantly, the candidates covered a large space of physicochemical properties, with various sequence lengths, amino acid preferences, stickiness, hydrophobicity, and net charge, all of which may impact phase separation propensity and IDR-IDR interaction specificity. We compared the sequence diversity of the selected sequences to that of all IDRs across the human proteome (**Methods**). We visualized the diversity of the IDR properties on a two-dimensional space using a uniform manifold approximation and projection (UMAP)^48^ (**Figure S2**). The selected IDRs were scattered across the map, highlighting that they cover a wide range of sequence characteristics on the IDR spectrum.

The nineteen IDR candidates were initially tested as monomeric chains fused to Venus and were expressed *in vivo*. However, only RBM14 and NUP98 formed condensates, with 12.5% and 25.2% of the cells identified with condensates, respectively (**Figure S3**). Only NUP98 fulfilled our criteria for determining phase separation as a monomeric construct (p=25.2%, s=0.07).

To ascertain the phase separation behavior was not heavily influenced by the fluorescent reporter or scaffold, we tested the candidate IDRs using two constructs: *micDROP*-green, and *micDROP*-red, which consists of the same overall architecture, but employs a different Dps-like scaffold from *S. solfataricus* (PDB code 2CLB^50^) and a red fluorescent protein (mScarlet^51^, **Methods**).

All nineteen candidates were tested in *micDROP*-red, and fifteen were additionally tested in *micDROP*-green. Thirteen candidates expressed well in both systems, and their phase separation behavior was consistent. Of those, seven did not meet our criteria for phase separation in both systems, while six (FUS, hnRNPA1, HspB8, RBM14, TAF15, and TDP43) exhibited a measurable C_sat_ (**Figure 2E, Figure S4, Figure S5**). The measured C_sat_ values were comparable between constructs (*R*=0.76, **Figure S2**), confirming that IDRs behaved similarly in both systems. Four additional candidates, Ddx4, DYRK3, ERα and UBQ2, showed phase separation propensity in *micDROP*-red but were not tested in *micDROP*-green (**Figure S5**).

The candidates not identified as phase separating fell into three categories. The first were lowly expressed (median intensity <100A.U.), and included CycT1, Ddx3x, and NUP98. Such low expression might be related to the toxicity and degradation of these particular constructs. Indeed, overexpression of disordered regions has been described as being frequently toxic^52^. The second category consisted of candidates forming condensates that did not fulfill our criteria for determining phase separation, either because <20% of the cells contained condensates, or because the dilute phase concentration did not appear stable among cells containing a condensate (s>0.4). This category included nuclear proteins (GATA3, NPM1 and HP1α) and might reflect incompatibility with the cytosolic localization we imposed, or could be caused by interactions of the system with other cellular components. The final category consisted of candidates that did not form condensates, even at high expression levels. These included our negative control (G3BP1) as well as Syn1. Several reasons may explain Syn1’s discrepancy. These include the solution conditions for Syn1’s phase separation *in vitro* (e.g., the requirement for PEG^53^), which may not be reflected *in vivo*, or the fusion of Syn1 to *micDROP*, which could inhibit its phase separation.

In summary, while nuclear and membrane-bound (NUP98, Nephrin) proteins were not suited for constitutive expression in the yeast cytosol, the majority of the remaining IDRs expressed well and were amenable to characterization with *micDROP*. In total, ten IDR candidates were identified as phase-separating and showed a range of C_sat_ values spanning two orders of magnitude. Such comparative data on a range of IDRs enabled investigating whether certain IDR properties were associated with self-recognition and phase separation propensity. To identify such properties, we compared C_sat_ values of IDRs to their respective features gathered for generating the UMAP. The strongest association appeared with the total stickiness of IDRs (*R*=-0.72, **Figure 2F, Figure S2**). This feature was previously identified as predictive of client recruitment into condensates^31^, which together with these observations implies that similar molecular interactions drive client-scaffold and scaffold-scaffold interactions.

### Average particle valency modulates IDR saturation concentration

Multivalent interactions are a key driving force underlying phase separation^37,54–56^. The avidity originating in multivalence can increase affinity and enables self-assembly while maintaining transient and dynamic interactions of individual sites, thus promoting a liquid-like state. Conversely, strong and stable interactions may not benefit from multivalency as much, as even a small number of interaction sites (e.g., four as opposed to twelve) could still promote the assembly of a network of physical interactions. In this case, however, such an assembly would not be dynamic and liquid-like.

We devised a simple strategy to modulate the valence of *micDROP*, guided by two primary motivations. First, we sought to test whether condensates with decreased valence were associated with increased C_sat_. Such correlation would indicate that *micDROP* was behaving in accordance with classic theories of phase separation, where a lower molecular valence, due to fewer interaction points, is typically associated with a higher C_sat_ required to induce phase separation. Conversely, we reasoned that constructs driving the formation of aggregates (e.g., due to misfolding or mis-assembly) would not respond to the valence modulation strategy. Secondly, we were motivated to characterize the impact of valency changes across different IDR systems to identify general behaviors or invariant characteristics. To modulate the average valency of the IDRs presented, we co-expressed a *micDROP*-red subunit with a ‘valence modulation’ subunit, composed of the same Dps scaffold fused to a different fluorescent reporter (Venus) with no IDR (**Figure 3A**). Upon co-expression, both subunits could co-assemble and form hybrid complexes, thereby modulating the number of IDRs presented per assembled particle. We harnessed the natural expression variability inherent to plasmids, whereby cells expressing both subunits on independent plasmids sampled an array of average particle valencies, which were estimated from the GFP and RFP fluorescence intensities in individual cells. For each strain expressing a specific IDR, the imaged cells were binned into five quintiles based on their GFP intensity, corresponding to the ‘valency modulator’ concentration (**Figure 3B, 3C**). The percentage of cells containing condensates in each quintile and their relative C_sat_ were then calculated as previously specified (**Figure 3D**).

**Figure 3.**
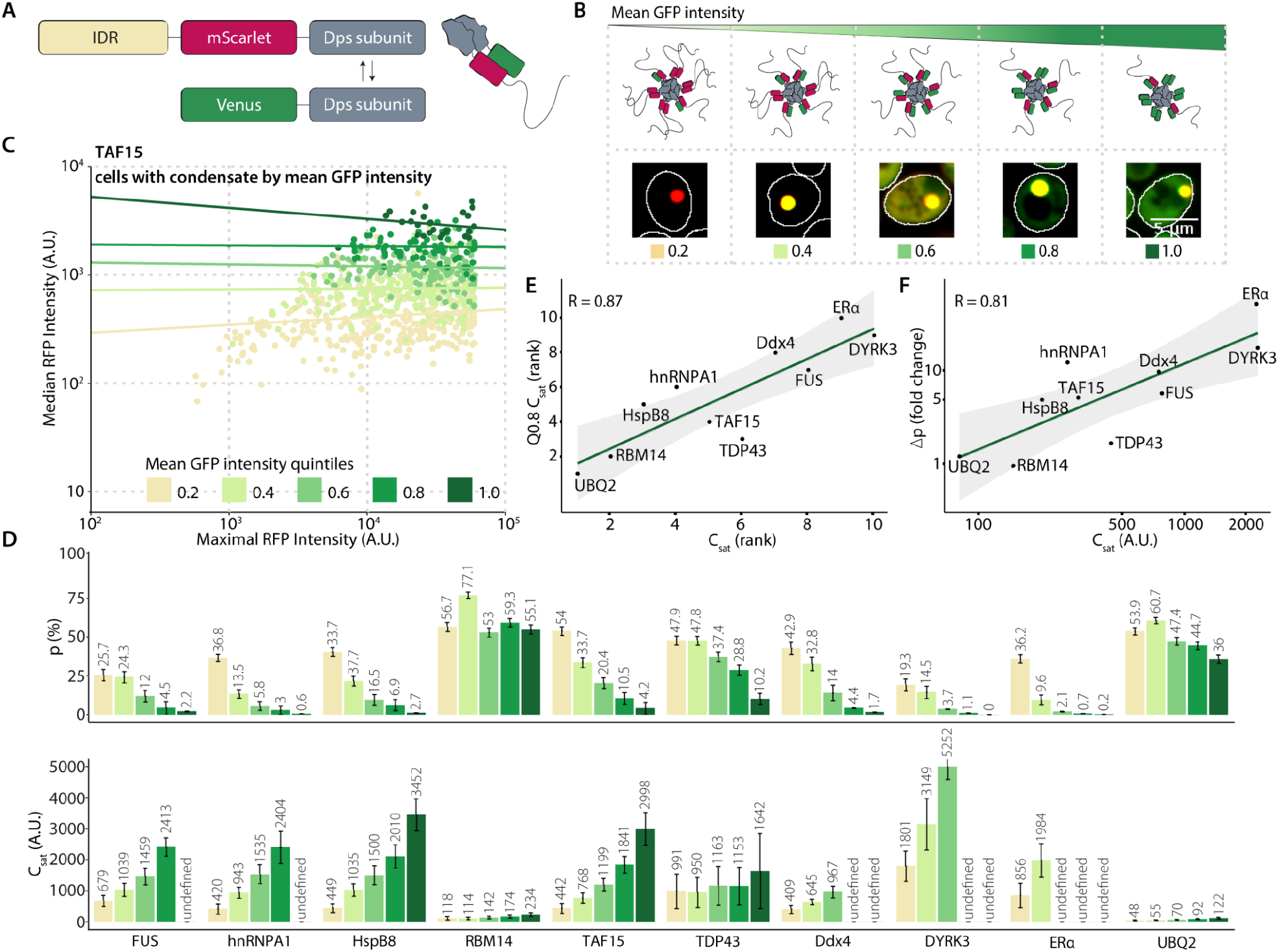
Average particle valency modulates IDR C_sat_. **A**. *micDROP*-red is co-expressed with a valency modulator consisting of the same Dps scaffold subunit fused to Venus and lacking an IDR. **B**. Co-expression of both constructs *in vivo* results in their co-assembly into hybrid scaffolds, with average IDR valency depending on their relative stoichiometries. Cells containing condensates that co-express *micDROP-red* with the valency modulator are binned into five quintiles by GFP intensity. Cells in the first quintile exhibit the lowest GFP levels and highest average particle valency. Conversely, cells in the fifth quintile exhibit the highest GFP levels and lowest valency. **C**. C_sat_ measurements of TAF15 in *micDROP*-red with a valency modulator. Each dot represents a cell containing a condensate, and is colored by quintiles of GFP intensity. The coordinates of each cell are the maximal (x-axis) and median (y-axis) intensity of RFP fluorescence. Decreased particle valency (high quintile) is associated with increased C_sat_. **D**. Barplots showing the percentage (*p*) of condensate-containing cells (top) and C_sat_ (bottom) per quintile of GFP expression, for ten IDR constructs. **E**. Rank-correlation of C_sat_ measurements of the first against fourth quintile. Missing values were ranked by data from other quintiles (Ddx4 < DYRK3 < ERα). **F**. Correlation of C_sat_ measurements against the fold-change in the percentage of condensate-containing cells between the first and fourth quintile. IDRs associated with high C_sat_ displayed a marked decrease in the percentage of condensate-containing cells, with the notable exception of TDP43, whereas IDRs with a low C_sat_ did not.

Of the ten IDRs previously identified as undergoing phase separation in *micDROP*-red, seven displayed a marked sensitivity to valency changes. FUS, hnRNPA1, HspB8, TAF15, Ddx4, DYRK3 and ERα showed a 5-fold or greater decrease in the percentage of condensate-containing cells between the first and fourth quintiles (**Figure 3D**). A similar effect was observed for their C_sat_ values, which increased 3 to 5-fold for FUS, hnRNPA1, HspB8, and TAF15, or increased to a point where they could not be detected for Ddx4, DYRK3, and ERα.

Interestingly, three IDR constructs, RBM14, TDP43, and UBQ2, exhibited decreased sensitivity to valency changes, with either no change, 1.6 or 1.2-fold decrease observed in the percentage of cells with condensates, and 1.5, 1.2 or 1.9-fold increase in C_sat_ between the first and fourth quintiles, respectively. We carried out the mirror experiment using *micDROP-*green (**Figure S6**), which confirmed the decreased sensitivity to effective valency changes for RBM14 and TDP43. UBQ2 was not tested in *micDROP*-green.

Overall, valency modulation preserved the relative ranking of C_sat_ across the different constructs: those exhibiting the lowest C_sat_ at high valency also showed the lowest C_sat_ when valency decreased (**Figure 3E**). Constructs exhibiting the lowest initial C_sat_ were also least affected in their propensity to form condensates, as reflected by the fold change in the percentage of cells containing condensates (**Figure 3F**). It is possible that stronger homotypic interactions in these constructs compensate for the change in valency in the concentration regimes of our experiments. Interestingly, however, we noted that TDP43 was an outlier in that respect. Despite having a measured C_sat_ considerably higher than that of RBM14 and UBQ2 (and also higher than those of HspB8, TAF15, and hnRNPA1), TDP43 showed a lack of sensitivity to valency changes. This observation appears at odds with the notion of strong IDR-IDR interactions and may be related to a liquid-to-solid maturation previously reported *in vivo* and *in vitro*^*57–59*^.

### TDP43 and UBQ2 localize into discrete condensates while other IDRs exhibit pronounced promiscuity

Our experiments yielded ten IDR candidates that underwent phase separation in *micDROP*-red with a wide range of C_sat_ and sensitivity to valency changes. We next sought to determine whether these IDRs were capable of forming discrete condensates through sequence-specific interactions. To this aim, we first validated that homologous scaffolds from *micDROP*-green and *micDROP*-red did not co-assemble (**Figure 4A**). We purified each scaffold in two forms: one untagged and another fused to a GFP tag. The untagged protein was incubated overnight with each of the GFP-tagged constructs, and the resulting particle size distribution was measured by mass photometry (**Methods**). Co-assembly was observed when the non-tagged scaffold was incubated with its identical GFP-tagged counterpart, but no co-assembly was detected between pairs of homologs (**Figure 4B, Figure S8**). In addition to these *in vitro* experiments, we ascertained that both homologs did not co-assemble *in vivo*. We co-expressed both scaffolds in cells, with the *micDROP*-green subunit fused to hnRNPA1 IDR, and the *micDROP*-red subunit without an IDR.

**Figure 4.**
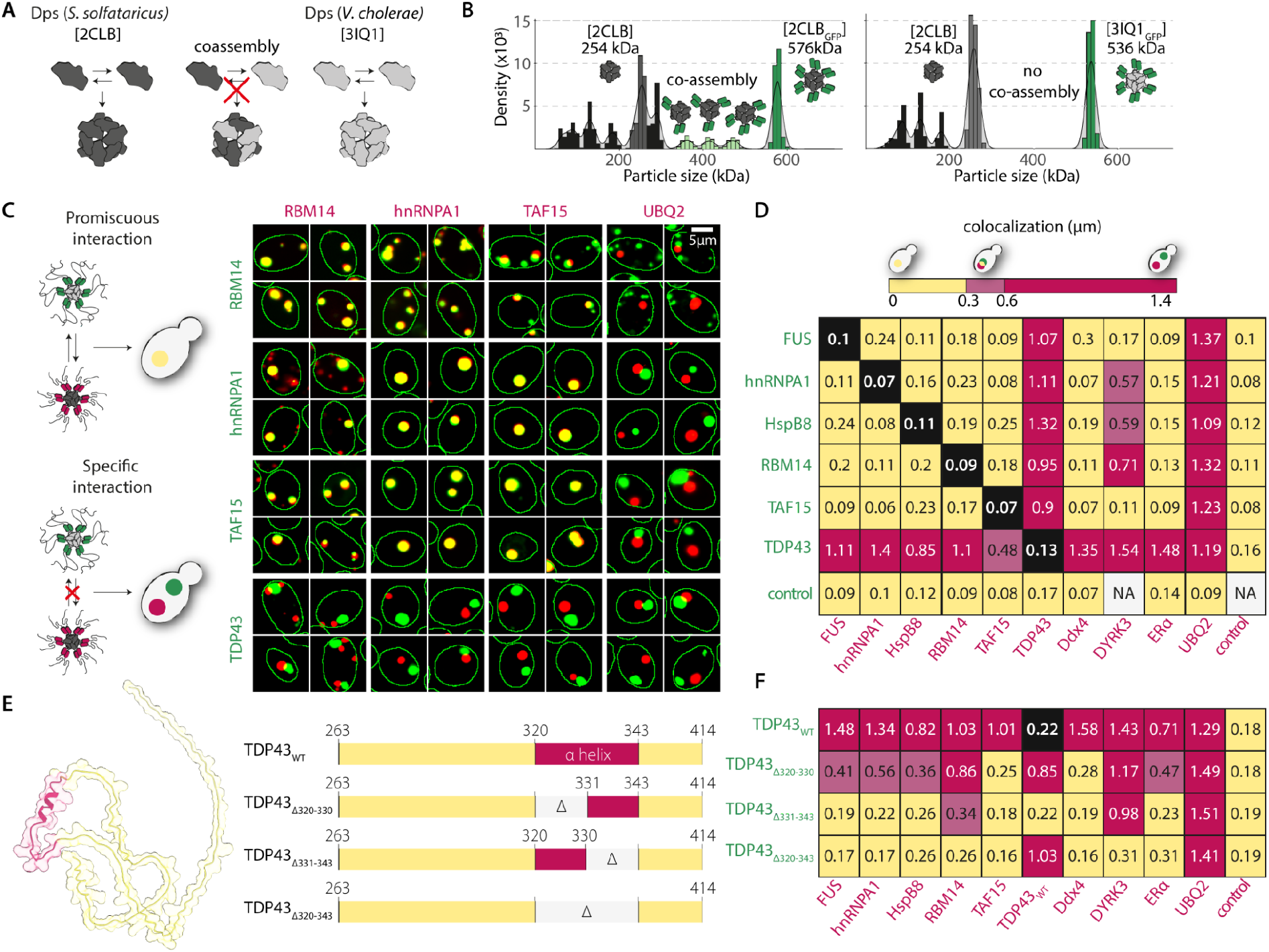
Most IDRs display high promiscuity while TDP43 forms discrete condensates through an α-helical segment. **A**. Probing IDR-IDR interaction specificity with *micDROP* necessitates two homologous scaffolds that do not co-assemble. We employed scaffolds from *S. solfataricus* (dark gray) and *V. cholerae* (light gray). **B**. We expressed and purified the scaffolds either tagged with GFP or untagged, and subsequently incubated both variants overnight to test their co-assembly by mass photometry. We observed hybrid complexes (left), but did not observe such co-assembly when incubating the homologous scaffolds (right). **C**. Schematic representation of specific vs. promiscuous IDR-IDR interactions (left). IDRs interacting promiscuously form condensates where both constructs colocalize, visible as a yellow puncta. Conversely, IDRs that interact in a self-specific manner yield spatially distinct condensates, as seen for TDP43 or UBQ2 in fluorescence microscopy images co-expressing micDROP-green and micDROP-red (right). **D**. Colocalization matrix showing the median distance (μm) between condensates formed by *micDROP*-green and *micDROP*-red. **E**. Cartoon rendering of the TDP43 disordered region, based on a model generated with AlphaFold2 (left). TDP43 contains a transient α-helical region (red) predicted to undergo disorder-to-order transition upon homotypic binding^68^. Three mutant constructs (Δ320-330, Δ331-343, and Δ320-343) were designed to assess whether partial or complete deletion of the α helix impacted TDP43’s specificity (left). **F**. Median distance (μm) between condensates of TDP43 variants and condensates formed by other IDRs. The variants Δ331-343 and Δ320-343 lost their homotypic specificity and colocalized with all IDRs, with the exception of UBQ2. The variant Δ320-330 showed intermediate specificity.

We did not detect the *micDROP*-red subunits enriched in condensates, which also indicated a lack of co-assembly (**Figure S8**). The mirror experiment, with the *micDROP*-red subunit fused to hnRNPA1 and the *micDROP*-green subunit with an identical scaffold and no IDR, also demonstrated a lack of co-assembly.

A matrix of strains was created wherein each strain co-expressed *micDROP*-green and *micDROP*-red fused to respective IDRs (**Figure 4C**). Considering cells in which condensates were identified both for *micDROP*-green and *micDROP*-red, we calculated the distance between the centers of the two nearest condensates. If the distance was smaller than 0.3μm (∼3 pixels), we inferred that the two IDRs colocalized. We decided on this distance because pairs harboring the same IDR consistently showed distances below this value. Based on this cutoff, we determined that eight of the ten IDRs displayed pronounced promiscuity and colocalized indiscriminately (**Figure 4D**); these correspond to FUS, hnRNPA1, HspB8, RBM14, TAF15, Ddx4, DYRK3 and ERα.

Among these constructs, the pair with the greatest similarity in sequence features was FUS and TAF15, as seen in the UMAP (**Figure S2**). Both are members of the FET (FUS, EWSR1, TAF15) family, similarly structured RNA-binding proteins (RBPs) that undergo phase separation through a Prion-like domain (PrLD) harboring Tyr and Arg ‘stickers’^22,60^. As such, it could be expected that they would colocalize indiscriminately^31^, and their partitioning into different biomolecular condensates is likely regulated by cellular localization signals^61^. hnRNPA1 and RBM14 are also RBPs undergoing phase separation through PrLDs, though their disordered regions display greater diversity from those of FUS and TAF15, primarily in length, stickiness, and net charge, as seen in the UMAP. Despite this increased diversity, hnRNPA1 and RBM14 colocalized with FUS and TAF15 and with each other, showing that favorable interactions between these IDRs were permissive. Interestingly, RBM14 previously showed a low C_sat_ and a marked insensitivity to valency changes (**Figure 3D**). Nevertheless, its lack of specificity implies that strong homotypic affinity is insufficient to form discrete condensates that do not intermix.

The FUS-HspB8 pair is known to interact, with HspB8’s folded α-crystallin domain preventing FUS maturation by chaperoning its RNA-binding domain^62,63^, which is distinct from the IDR region. Interestingly, our results imply that interactions between the IDR regions of these two proteins are also favorable.

Among the co-localizing group of IDRs, the two furthest away from the RBP group (FUS, hnRNPA1, TAF15, RBM14) on the UMAP were Ddx4 and DYRK3, indicating they had the greatest diversity in their sequence features. Ddx4’s phase separation is driven by electrostatic interactions, with repeating residue blocks of alternating net charge, but also contains an over-representation of Phe-Gly, Gly-Phe, Arg-Gly, and Gly-Arg motifs within the positively charged blocks^12^. Our results indicate that these conserved motifs are sufficient to promote favorable interactions with PrLDs even in cases of strong homotypic affinity such as RBM14. The dual-specificity kinase DYRK3 partitions into stress granules via its intrinsically disordered N-terminal domain and its kinase activity is required for the dissolution of stress granules, possibly by phosphorylating multiple RBPs^64^. The IDR construct (not including the kinase domain) colocalized with FUS and TAF15, but displayed only intermediate specificity when co-expressed with hnRNPA1, RBM14 and HspB8. This behavior was unexpected, as it implies some unique mechanism or conformation of the DYRK3 IDR, which can interact differently with members of the ‘promiscuous’ group.

The IDRs of TDP43 and UBQ2, which were located closer to the RBP group (TAF15, FUS, hnRNPA1 and RBM14) on the UMAP compared to DYRK3 or Ddx4, nevertheless formed discrete condensates regardless of their IDR partner. Such behavior implies these IDRs contain additional features driving their exclusive self-recognition. Upon examining the two sequences, we observed that both TDP43 and UBQ2 contain short structured segments in their disordered sequence, which might account for their specificity. First, UBQ2 has a ubiquitin-associated domain (UBA) at its C-terminus, which recognizes polyubiquitin chains on marked proteins^65,66^. In the future, *micDROP* could serve to test mutant sequences of UBQ2 and examine the effect of the UBA segment on UBQ2’s phase separation propensity and self-recognition specificity. Second, TDP43 has an evolutionarily conserved transient α-helical segment known to modulate its phase separation propensity^67,68^. This segment can undergo disorder-to-order transition upon homotypic binding, which we hypothesized could account for TDP43’s specificity.

### A transient α-helix in TDP43’s disordered region governs its self-specific condensation

The disordered region of TDP43 is a PrLD that lacks extensive RGG repeats compared to other IDRs from RNA-binding proteins such as FUS or RBM14. However, it contains a conserved region (**Figure 4E**) that forms a transient α helix that promotes homotypic interactions^57^. To test whether this region mediates the specific self-recognition of TDP43, we generated three mutant sequences with either partial (Δ320-330 and Δ331-343) or complete (Δ320-343) deletions of this segment and tested their colocalization with other IDRs using *micDROP*. Variant Δ320-330 showed intermediate specificity with other IDRs and surprisingly, lost the ability to colocalize with TDP43 WT. Variant Δ331-343 lost all specificity and colocalized promiscuously with all IDRs, including TDP43 WT. Variant Δ320-343 also colocalized promiscuously with all other IDRs, but lost the ability to colocalize with TDP43 WT (**Figure 4F, Figure S8**). We conclude that TDP43’s specificity is governed by the α helical segment, and hypothesize that the 320-330 region modulates homotypic recognition while the 331-343 segment is likely required to stabilize the interaction.

## CONCLUSIONS

We developed a synthetic system called *micDROP* that proved powerful in studying the potential of IDRs to drive intracellular phase separation. Characterizing interaction properties of IDRs is notoriously difficult, particularly *in vivo*, owing to their conformational flexibility. Nevertheless, we showed that *micDROP* captured the phase separation propensity of various IDRs *in vivo* by quantifying their saturation concentrations. The simplicity of *micDROP*’s design enabled us to analyze a large number of IDRs, which revealed a correlation between an IDR’s C_sat_ and its total sequence stickiness. We also showed that the effective valency of *micDROP* could be easily modulated, which revealed both expected phase separation behavior for most IDRs, and a surprising one for TDP43. The characterization of two homologous scaffolds that did not co-assemble made *micDROP* amenable to studying IDR self-recognition specificity. This feature is unique, as recent work has been employing different scaffold geometries to this effect^69^, which adds confounding factors absent from *micDROP*.

Equally important, the modular nature of the system means it is readily generalizable to other IDR sequences of interest. Our findings indicate that IDRs capable of undergoing phase separation are generally promiscuous owing to their interactions through nonspecific sticky patches^31^. IDRs that formed discrete condensates contained structured segments, reinforcing the notion that fully disordered and flexible regions undergoing phase separation are inherently non-specific. These results imply that regulating the formation and composition of biomolecular condensates should generally require complex networks of IDRs functioning hand-in-hand with structured partners or latent structural elements encoded in their sequences.

## METHODS

### Ferritin scaffold candidate selection

The list of candidate ferritins was obtained from known structures found in the Protein Data Bank (PDB)^70^, which we mined based on the 3DComplex database^71^. Specifically, we curated a list of 24-mer ferritin and 12-mer ferritin-like proteins non-redundant at a level of 40% sequence identity. Their UniProt sequences were aligned using Clustal Omega MSA tool^72^, and eight final candidates were selected (**Table S2**).

### Ferritin scaffold cloning, expression and purification in *E. coli*

All UniProt sequences were optimized for bacterial expression, and synthesized into two pET T7 RNA polymerase-driven transcription vectors (Twist Bioscience) with the following sequence features:

> **Plasmid 1:** [6xHIS] [TEV (tobacco etch virus)] [Ferritin scaffold] [Ampicillin resistance cassette]
>
> **Plasmid 2:** [6xHIS] [TEV] [super-folder Green Fluorescent Protein (sfGFP)] [linker] [Ferritin scaffold] [Kanamycin resistance cassette]

Plasmids were transformed into E. coli BL21-CodonPlus (DE3)-RIPL cells. Transformant cells were grown in LB media at 37°C with appropriate antibiotics to OD_600_ ∼0.8. Expression was induced by the addition of 1mM isopropyl b-D-1-thiogalactopyranoside (IPTG) and allowed to proceed overnight (∼7h) at 25°C. Cells were harvested using French press and centrifugation. Each protein was purified on a 5ml nickel nitrilotriacetic acid (Ni^2+^-NTA) resin (GE Healthcare). The column was washed stepwise with 20, 50, 100 and 250mM imidazole. Each protein was eluted with 500mM imidazole, and the N-terminal 6xHIS tag was removed by TEV protease cleavage. Each protein was then purified on a HiPrep™ 16/60 Sephacryl S-500 HR gel filtration column (GE Healthcare) equilibrated with 50mM HEPES (pH 7.5), 50mM KCl and 1mM dithiothreitol (DTT).

### Assessing co-assembly of ferritin scaffolds using mass photometry

Purified protein samples were prepared for imaging by mixing each non-tagged scaffold with all GFP-tagged constructs at a 1:1 concentration ratio, followed by overnight incubation at 10°C. Immediately prior to mass photometry measurements, samples were diluted using stock buffer (50mM HEPES (pH 7.5), 50mM KCl and 1mM dithiothreitol (DTT)). Measurements were performed using OneMP Refeyn Ltd. mass photometer. To find focus, a fresh stock buffer was first flown into the chamber, the focal position was identified and secured in place with an autofocus system based on total internal reflection for the entire measurement period. Typical working concentrations of protein mixtures were ∼25–50nM, and each sample was measured in a new flow-chamber. For each acquisition, ∼15μL of diluted protein mix was introduced into the flow-chamber and, following autofocus stabilization, movies of 60-90s were recorded. Each sample was measured three times independently. The molecular weight was obtained by contrast comparison with known mass standard calibrants measured on the same day.

### UMAP and IDR candidate selection

A uniform manifold approximation and projection (UMAP) map was generated for the entire human disordered proteome (all protein segments of ≥35 residues predicted to be disordered according to consensus predictions reported in MobiDB ^73^ (**Table S1, Figure S2**). The number of neighbors was set to 20 or 40 to assess clustering stability across this range. Variables taken into account when generating the UMAP included:

- Amino acid enrichment compared to entire human disordered proteome
- Frequencies and over-representation of amino acids
- Net charge
- pI
- Hydrophobicity^74^
- Stickiness^31^
- AGGRESCAN values^75^

The entries of human IDRs listed in the PhaSePro database^45^ and that we selected were highlighted onto the UMAP. The final candidate list (**Table 1**) was generated by considering UMAP distances, sequence length, overall net charge, and participation in functionally distinct biomolecular condensates.

### IDR cloning and expression in *S. cerevisiae*

UniProt sequences were optimized for expression in *S. cerevisiae*. Fifteen candidates were synthesized (Twist Bioscience) in two custom expression vectors containing the following features:

> ***micDROP*-green:** [TDH3 promoter] [Nuclear export signal (NES)] [IDR] [linker] [YFP Venus] [linker] [Ferritin scaffold] [CYC terminator] [Hygromycin resistance cassette]
>
> ***micDROP*-red:** [TDH3 promoter] [NES] [IDR] [linker] [RFP mScarlet] [linker] [Ferritin scaffold] [CYC terminator] [Nourseothricin resistance cassette]

Six additional candidates were synthesized only into *micDROP*-red. *micDROP*-red plasmids were transformed into *S. cerevisiae* BY4741 (MATa his3Δ1 leu2Δ0 met15Δ0 ura3Δ0). *micDROP*-green plasmids were transformed into *S. cerevisiae* BY4742 (MATα his3Δ1 leu2Δ0 lys2Δ0 ura3Δ0). Haploid strains were then mated on YPD agar plate and allowed to grow overnight (∼12h). Diploid selection was repeated twice on YPD agar plate with Hygromycin and Nourseothricin selections.

### Confocal fluorescence microscopy, image processing, and analysis

For imaging, 1 μl of saturated cell suspension was transferred to an optical plate with 29μl fresh SD media with appropriate antibiotics and grown at 30°C for 3h to logarithmic growth. Cells were imaged with an Olympus IX83 microscope coupled to a Yokogawa CSU-W1 spinning-disc confocal scanner with dual prime-BSI sCMOS cameras (Photometrix). The 16-bit images were acquired with three illumination schemes of 50 ms exposure each: one for brightfield, one for green, and one for red fluorescence. For green fluorescence, we excited the sample with a 488 nm laser (Coherent 150 mW) and collected light through a bandpass emission (Em) filter (525/50 nm, Chroma ET/m). For red fluorescence, we excited at 561 nm (Coherent 150 mW) and used a bandpass emission filter (609/54 nm, Chroma ET/m). One multiband dichroic mirror was used for the three illuminations. Imaging was performed with a ×60/1.42 numerical aperture (NA), oil-immersion objective (UPLSAPO60XO, Olympus).

Following acquisition, cells were identified, segmented and their fluorescent signal (median, average, minimum, maximum, 10^th^, 20^th^, …, 90^th^ percentile fluorescence) were identified using previously reported scripts in ImageJ^43^. Condensates were identified in each cell independently, in a multistep process: (1) calculation of the median fluorescence intensity of pixels in a given cell; and (2) identification of the largest region composed of pixels with an intensity twofold (GFP) or threefold (RFP) above the median. If such a region existed and it showed a circularity over 0.4, the condensate properties (intensity, size, etc), and its coordinates were recorded. Tabulated data from image analyses were loaded and analyzed further with custom scripts in R.

### Analysis of IDRs’ phase separation propensity *in vivo*

We noticed that very high concentrations of RFP gave a noticeable (1-2%) cross-talk signal in the GFP channel. Therefore, we applied a GFP channel brightness correction to account for this crosstalk. We subtracted 2% of RFP brightness values from the GFP values. Additionally, we discarded all cells where the GFP intensity was saturated (value above 65536) as no correction could be reliably applied for those.

Condensates detected during image analysis were deemed valid if the mean brightness within the condensate was significantly higher than background fluorescence (>150 arbitrary fluorescence units, AFUs) after the correction for crosstalk was applied. If a strain had >20 segmented cells with valid condensates and at least 20% of the segmented cells were identified as containing a condensate, we analyzed the propensity of the corresponding IDR to phase separate by analyzing the slope of points defined by maximal (x-axis) and median (y-axis) fluorescence intensities within each cell.

### Analysis of IDR colocalization *in vivo*

Colocalization calculations were done for IDR candidates previously identified as phase separating. For each strain, we ensured that at least 50 cells containing condensates in both fluorescent channels were used to calculate a co-localization score. We used the minimal Euclidean distance between condensates, based on the largest two condensates per channel. Thus, the minimum of (at most four) pairwise distances was recorded for each cell. Subsequently, the median of these distances gave the colocalization score for the corresponding strain.

### Generation of structure models

The structure models were generated with AlphaFold2 Multimer^41,76^ using a local implementation of Colabfold^77^.

## Supporting information

Supplemental_FigsS1-9_TableS2

## Data Availability

The code and data used to generate the UMAP is available at: https://github.com/benjamin-elusers/human_idr.

Plasmids and strains are available upon request.

## Authors contributions

AG and EDL designed and planned the research. AG carried out the experimental work and analyzed the microscopy data. AG and EDL interpreted the results. BD carried out the bioinformatics analyses on IDR regions to generate the UMAP. AG and EDL wrote the manuscript with input from BD.

## Acknowledgments

We thank Prof. Sam Safran of the Department of Chemical and Biological Physics at the Weizmann Institute of Science for thoughtful ideas and suggestions regarding phase separation theory, Prof. Michael Elbaum and Dr. Alexandra Tayar for their insightful feedback on experimental design and analysis, Prof. Rina Rosenzweig of the Department of Chemical and Structural Biology at the Weizmann Institute of Science for her advice on chaperone selection, Dr. Adar Sonn-Segev for her expertise with mass photometry, and past and current members of the Levy lab for their invaluable support and assistance. This work was supported by the European Research Council (ERC) under the European Union’s Horizon 2020 research and innovation program (grant agreement No. 819318), by the Israel Science Foundation (grant no. 1452/18), by a research grant from A.-M. Boucher, by research grants from the Estelle Funk Foundation, the Estate of Fannie Sherr, the Estate of Albert Delighter, the Merle S. Cahn Foundation, Mrs. Mildred S. Gosden, the Estate of Elizabeth Wachsman, the Arnold Bortman Family Foundation.

