## Supplemental_FigsS1-9_TableS2 for "Mapping interactions between disordered regions reveals promiscuity in biomolecular condensate formation"

### SUPPLEMENTARY MATERIAL

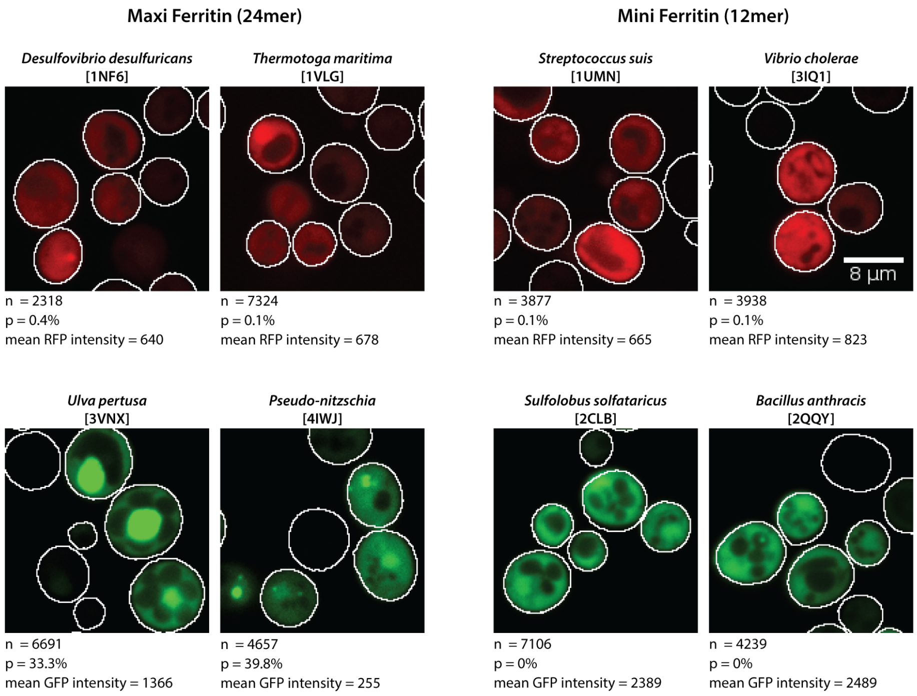


**Figure S1. Assessment of candidate scaffold for *in vivo* expression.** Fluorescence microscopy images of live cells expressing four 24-mer (left) and four 12-mer (right) candidate scaffolds tagged with RFP (mScarlet, top) or YFP (Venus, bottom). YFP-tagged 24-mer scaffolds frequently formed puncta (p) and were therefore discarded.

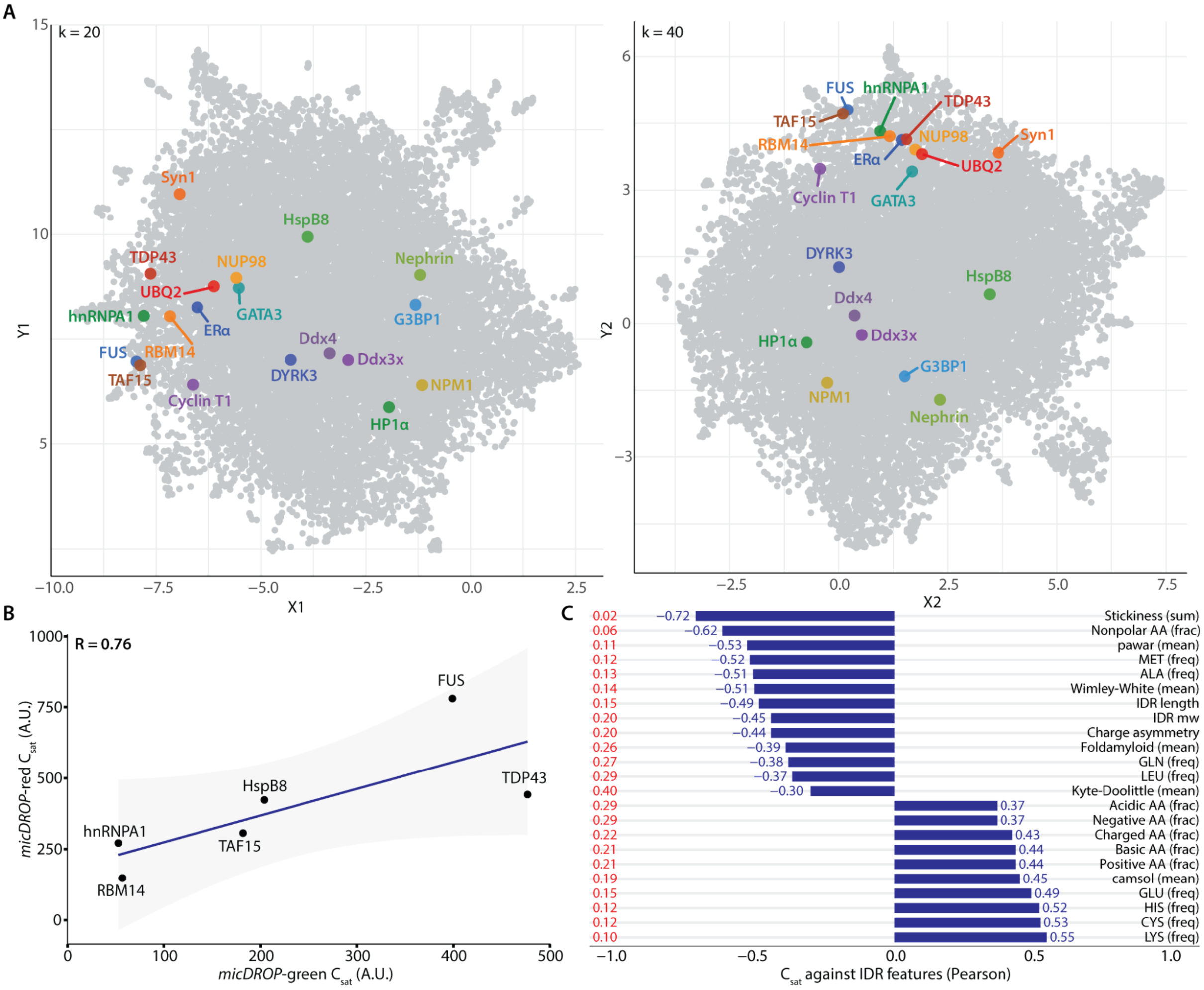


**Figure S2. Computational analysis of features of intrinsically-disordered regions undergoing phase separation. A.** UMAP projecting the diversity of human IDR sequences in 2D. Maps were generated with either twenty (left) or forty neighbors (right). **B.** $C_{sat}$ values are comparable between *micDROP*-green and *micDROP*-red for different IDRs. **C.** Pearson correlation between measured $C_{sat}$ values in *micDROP*-red and features considered in the UMAP. We find that phase separation propensity shows the strongest association with the sum of sequence stickiness.

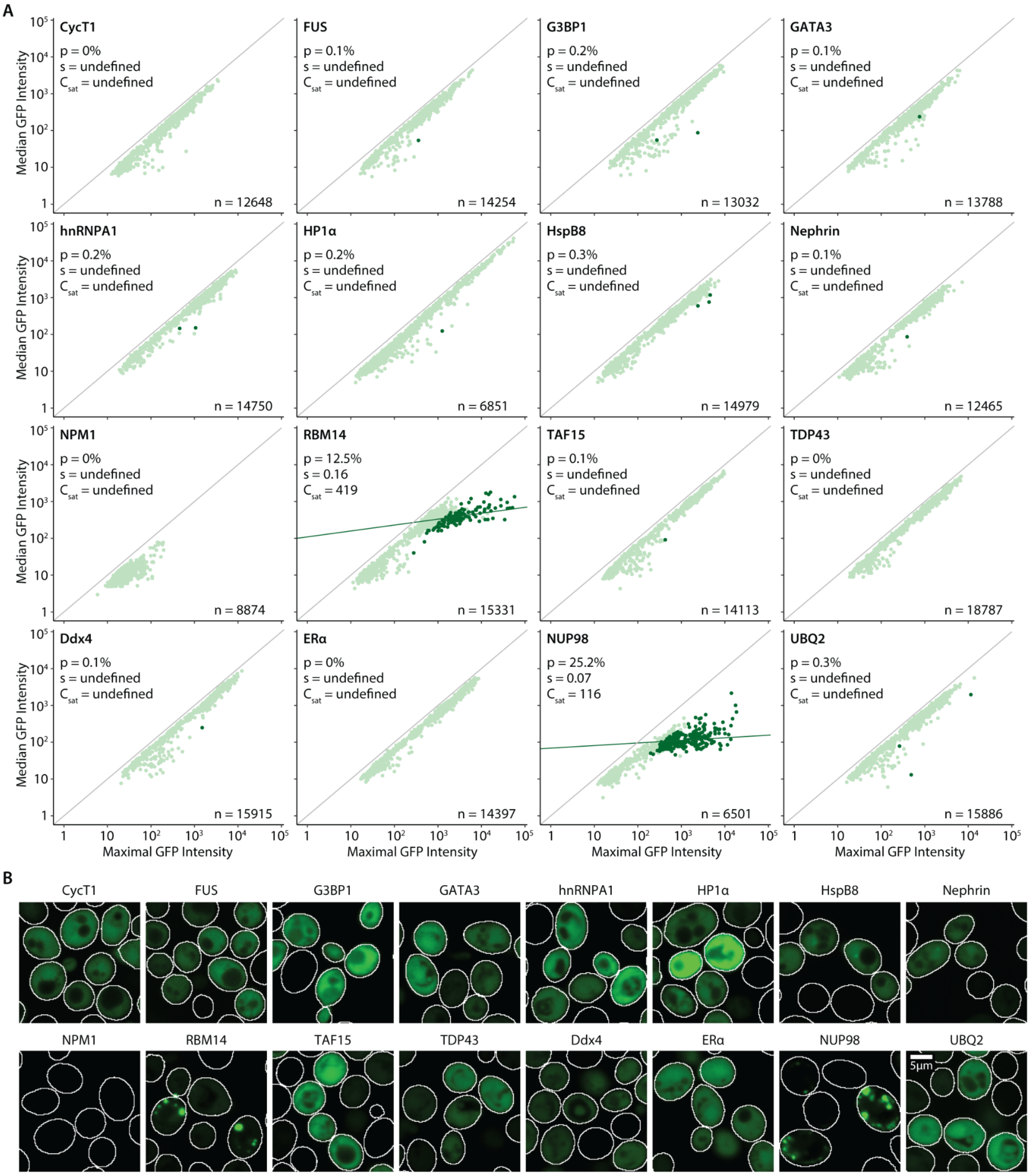


**Figure S3. $C_{sat}$ measurements of monomeric IDRs. A.** $C_{sat}$ measurements of candidate IDRs tagged with YFP (Venus) show that most IDRs do not form condensates *in vivo* when expressed as monomeric constructs, with the exception of NUP98 and RBM14. To facilitate comparison we show the same number of randomly sampled data points (n=1000) for each scatterplot. **B.** Fluorescence microscopy images of candidate IDRs expressed as monomeric constructs *in vivo*.

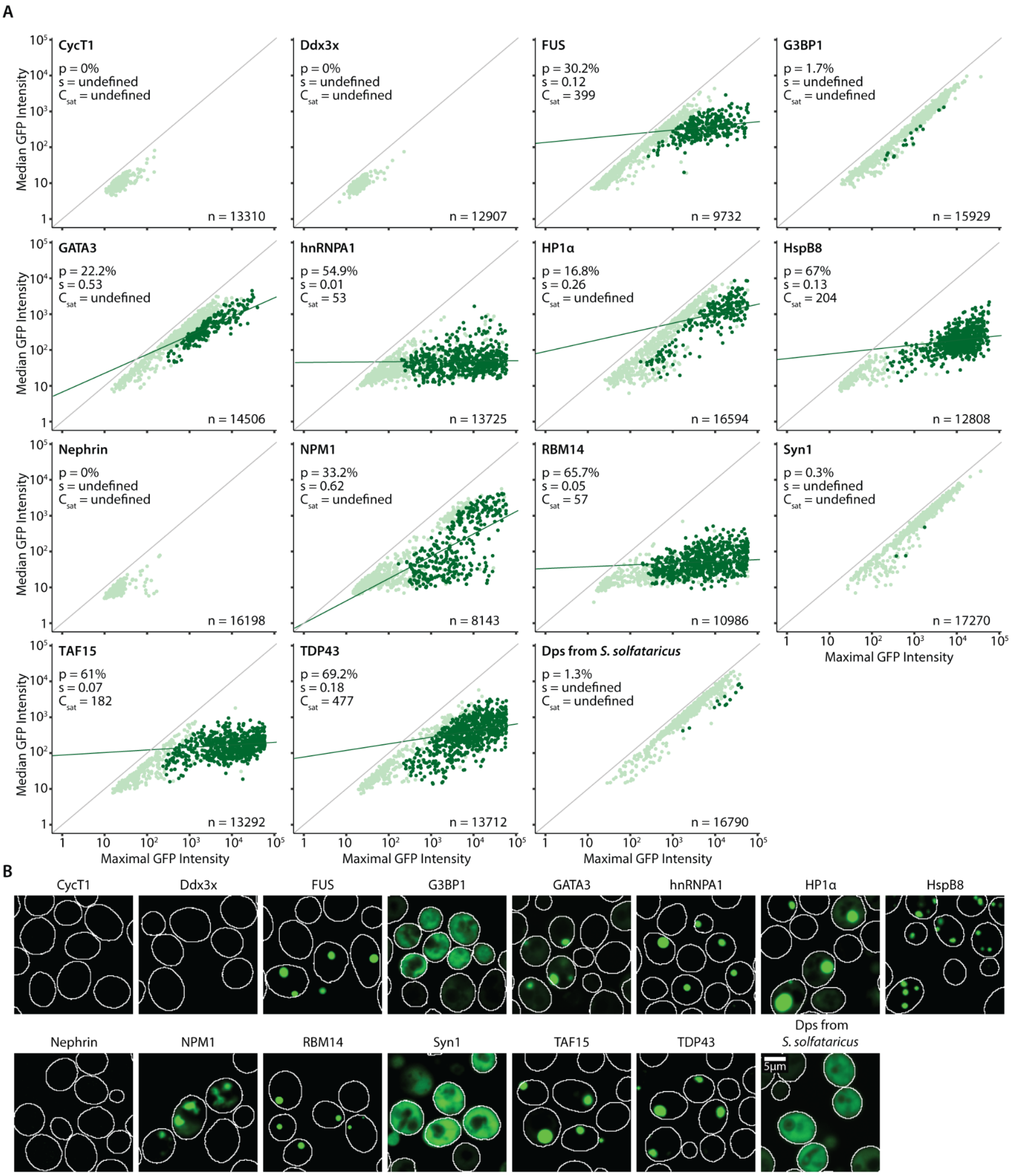


**Figure S4. Measuring $C_{sat}$ for IDRs expressed with *micDROP*-green. A.** $C_{sat}$ measurements of candidate IDRs fused to *micDROP*-green. To facilitate comparison we show the same number of randomly sampled data points (n=1000) for each scatterplot. In the absence of an IDR, *micDROP*-green does not form condensates *in vivo* (p=1.3%). **B.** Fluorescence microscopy images of candidate IDRs fused to *micDROP*-green *in vivo*.

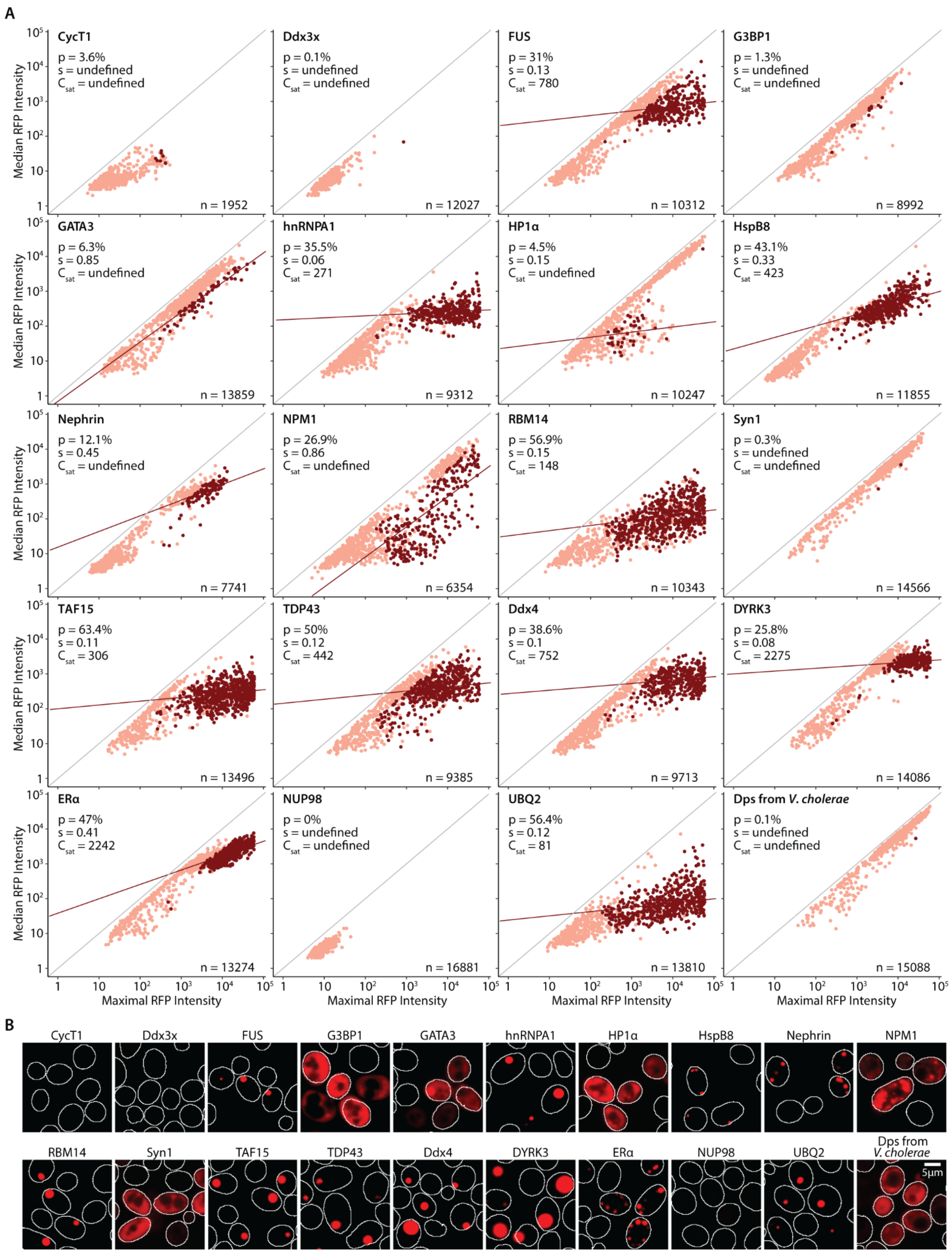


**Figure S5. Measuring $C_{sat}$ for IDRs expressed with *micDROP*-red. A.** $C_{sat}$ measurements of candidate IDRs fused to *micDROP*-red. To facilitate comparison we show the same number of randomly sampled data points (n=1000) for each scatterplot. In the absence of an IDR, *micDROP*-red does not form condensates *in vivo* (p=0.1%). **B.** Fluorescence microscopy images of candidate IDRs fused to *micDROP*-red *in vivo*.

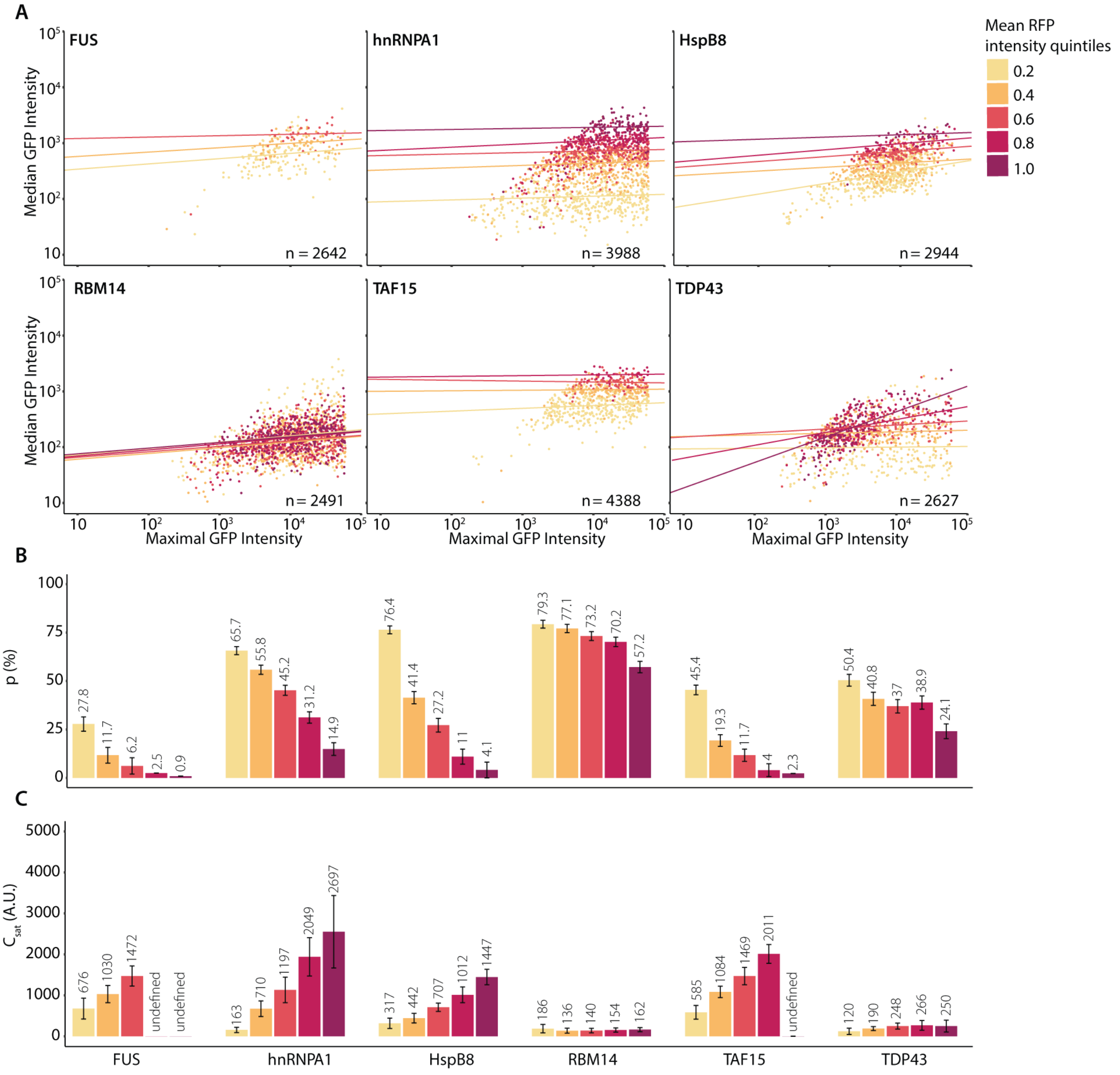


**Figure S6. Decreasing average particle valency increases $C_{sat}$ in *micDROP*-green A.** $C_{sat}$ measurements of six phase separating IDRs in *micDROP*-green co-expressed with a valency modulator labeled with a red fluorescent reporter (mScarlet). Each dot represents a cell containing a condensate and is colored by quintiles of RFP intensity. The coordinates of each cell are the maximal (x-axis) and median (y-axis) intensity of GFP fluorescence. Decreased particle valency is generally associated with increased $C_{sat}$. **B.** Barplots showing the percentage (p) of condensate-containing cells (top) and $C_{sat}$ (bottom) per quintile of valency modulator (RFP) expression.

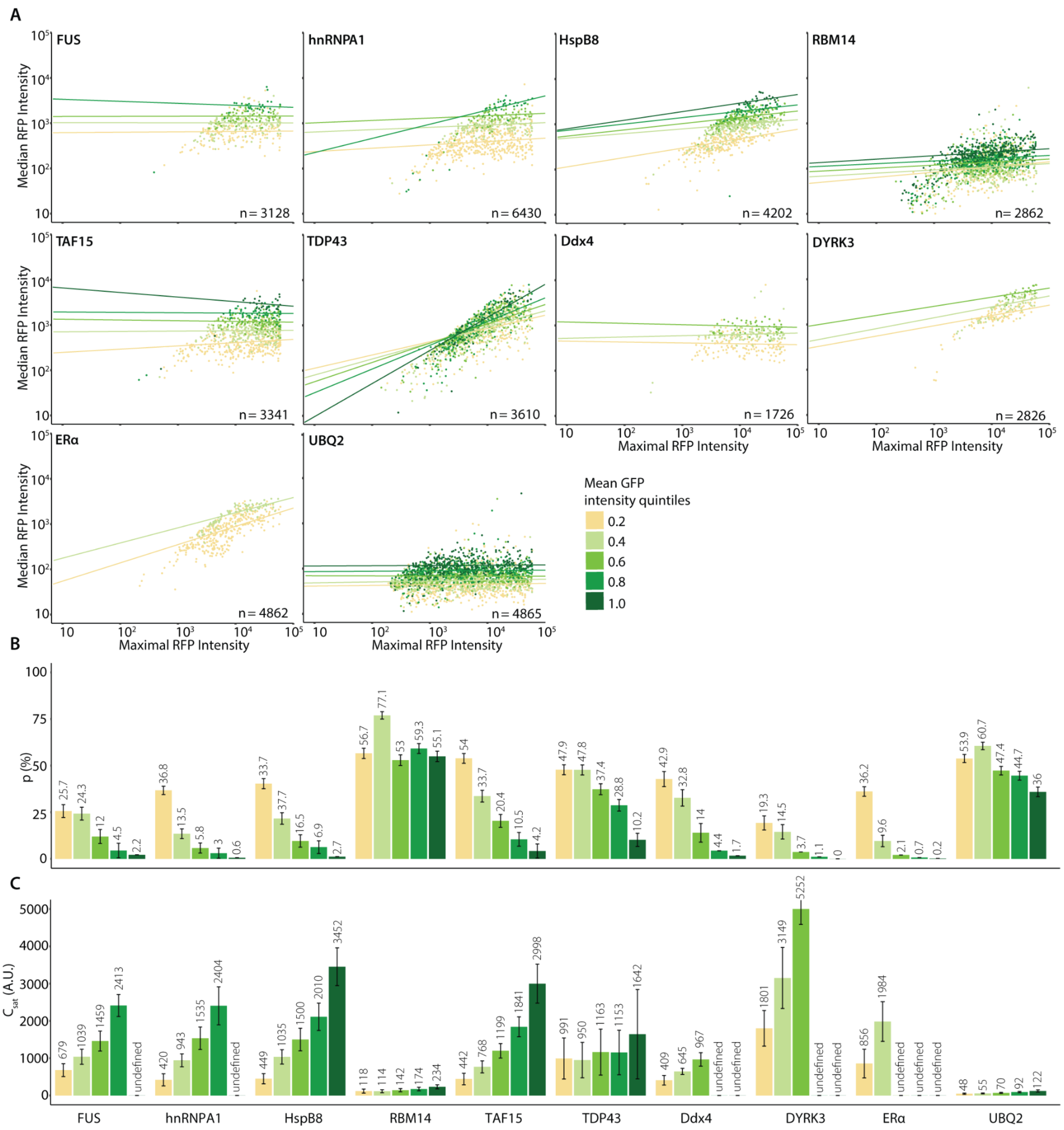


**Figure S7. Decreasing average particle valency increases $C_{sat}$ in *micDROP*-red A.** $C_{sat}$ measurements of ten phase separating IDRs in *micDROP*-red co-expressed with a valency modulator labeled with a yellow fluorescent reporter (Venus). Each dot represents a cell containing a condensate and is colored by quintiles of GFP intensity. The coordinates of each cell are the maximal (x-axis) and median (y-axis) intensity of RFP fluorescence. Decreased particle valency is generally associated with increased $C_{sat}$. **B.** Barplots showing the percentage (p) of condensate-containing cells (top) and $C_{sat}$ (bottom) per quintile of valency modulator (GFP) expression.

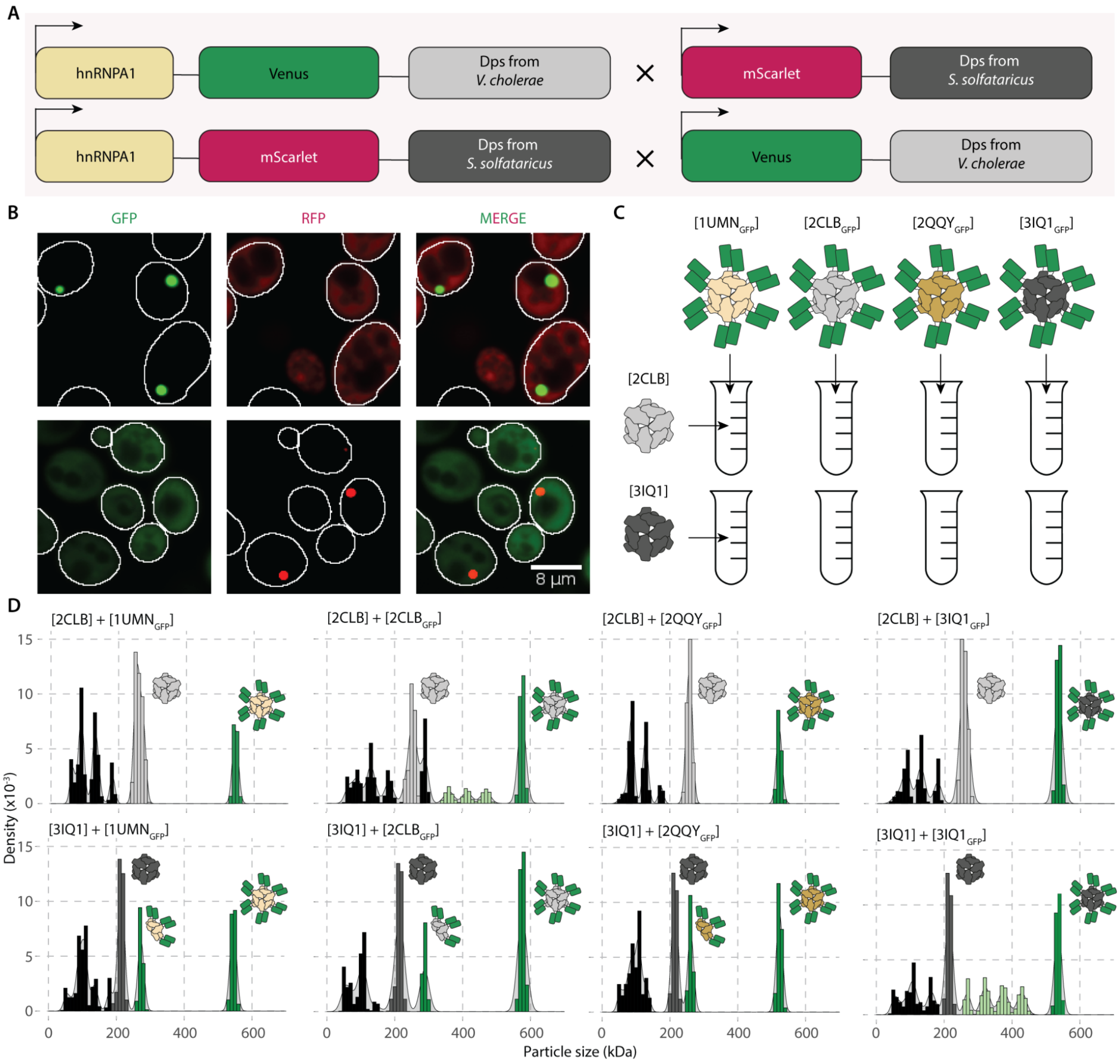


**Figure S8. Homologous Dps scaffolds do not co-assemble. A.** To test whether the homologous Dps scaffolds underwent co-assembly *in vivo*, *micDROP*-green fused to hnRNPA1 IDR was co-expressed with *micDROP*-red with no IDR (top), and *micDROP*-red fused to hnRNPA1 IDR is co-expressed with *micDROP*-green with no IDR (bottom). **B.** Fluorescence microscopy images of cells expressing two *micDROP* constructs. When hnRNPA1 is fused to *micDROP*-green (top) it forms condensates, while *micDROP*-red remains dispersed in the cytoplasm, and when hnRNPA1 is fused to *micDROP*-red (bottom) it forms condensates, while *micDROP*-green remains dispersed. **C.** Four homologous Dps scaffolds were purified and then incubated together overnight before measuring their particle size distribution by mass photometry. **D.** We observed hybrid complexes when the Dps scaffold was mixed with the same GFP-tagged subunit, but no co-assembly was observed when incubating the homologous scaffolds.

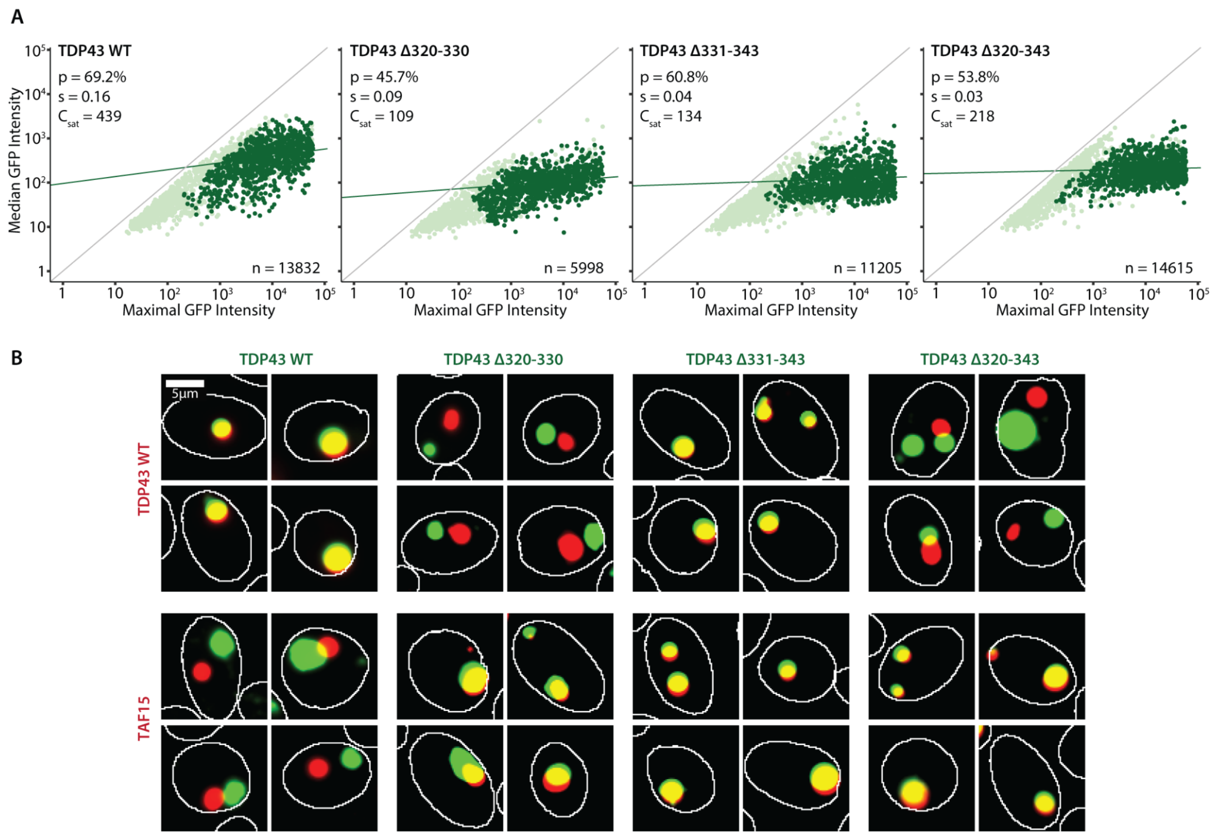


**Figure S9. Partial or complete deletion of the α helix segment in TDP43's disordered region does not inhibit phase separation but increases promiscuity. A.** $C_{sat}$ measurements of three TDP43 variants using *micDROP*-green reveal that partial or complete deletion of the α helix segment does not inhibit TDP43's phase separation, but does affect the observed $C_{sat}$. **B.** Fluorescence microscopy images of TDP43 WT and its three variants expressed with *micDROP*-red TDP43 WT (top), and with TAF15 (bottom).

#### Table S1. List of features for all disordered regions of the human proteome. [LINK]

**Table S2. Ferritin scaffold candidates**

| PDB Code | Organism | Organism Type | Valency | Ferritin Type | AA Sequence |
|---|---|---|---|---|---|
| 1NF6[78] | *Desulfovibrio desulfuricans* | Gram-negative sulfate-reducing bacteria | 24 | Bacterioferritin | MAGNREDRKAKVIEVLNKARAMELHAIHQYM NQHYSLDDMDYGELAANMKLIAIDEMRHAEN FAERIKELGGEPTTQKEGKVVTGQAVPVIYES DADQEDATIEAYSQFLKVCKEQGDIVTARLFE RIIEEEQAHLTYYENIGSHIKNLGDTYLAKIAGT PSSTGTASKGFVTATPAAE |
| 1VLG | *Thermotoga maritima* | Gram-negative, hyperthermophilic, anaerobic bacteria | 24 | Ferritin | MMVISEKVRKALNDQLNREIYSSYLYLSMATY FDAEGFKGFAHWMKKQAQEELTHAMKFYEYI YERGGRVELEAIEKPPSNWNGIKDAFEAALKH EEFVTQSIYNILELASEEKDHATVSFLKWFVDE QVEEEDQVREILDLLEKANGQMSVIFQLDRYL GQRE |
| 3VNX[79] | *Ulva pertusa* | Green algae | 24 | Ferritin | MLSASIKASTGATKAVGAGRLSHFQLRRQRG VSAHAAQEVTGMVFQPFSEVQGELSTVTQAP VTDSYARVEYHIECEAAINEQINIEYTISYVYHA LHSYFARDNVGLPGFAKFFKEASDEEREHAH MLMDYQTKRGGRVELKPLAAPEMEFANDDK GEALYAMELALSLEKLNFQKLQALQAIADKHK DAALCDFVEGGLLSEQVDAVKEHAVYVSQLR RVGKGVGVYLLDQELGEEEA |
| 4IWJ[80] | *Pseudo-nitzschia* | Diatom algae | 24 | Ferritin | MKSPFFFLSALALTLRDSSPSFATAFRLAVTRC ARQGIHAPSSSSSSSSSRCLVASASALAGPSEE LLDLFNRQVTQEFTASQVYLSASIWFDQNDW EGMAAYMLAESAEEREHGLGFVDFANKRNIPI ELQAVPAPVSCAEWSSPEDVWQSILELEQAN TRSLLNLAEAASTCHDFAVMAFLNPFHLQQVN EEDKIGSILAKVTDENRTPGLLRSLDVVSFLGP CLFRSV |
| 1UMN[81] | *Streptococcus suis* | Gram-positive bacteria | 12 | Dps-like | MMKQKYYQSPAEIASFSPRPSLADSKAVLNQ AVADLSVAHSILHQVHWYMRGRGFMIWHPKM DEYMEEIDGYLDEMSERLITLGGAPFSTLKEF SENSQLKEVLGDYNVTIEEQLARVVEVFRYLA ALFQKGFDVSDEEGDSVTNDIFNVAKASIEKHI WMLQAELGQAPKL |
| 2CLB[50] | *Sulfolobus solfataricus* | Thermophilic archaea | 12 | Dps-like | MQEKPQEPKVVGVEILEKSGLDIKKLVDKLVK ATAAEFTTYYYYTILRMHLTGMEGEGLKEIAED ARLEDRLHFELMTQRIYELGGGLPRDIRQLADI SACSDAYLPENWKDPKEILKVLLEAEQCAIRT WKEVCDMTYGKDPRTYDLAQRILQEEIEHEA WFLELLYGRPSGHFRRSSPGNAPYSKK |
| 2QQY | *Bacillus anthracis* | Gram-positive bacteria | 12 | Dps-like | MSHDVKELIEGLNEDLAGEYSAIIMYNHNAATV SGIYRQVLKPFFESEISDEQGHALYLAEKIKTL GGTPTTIPLRVKQAEDVREMLEYARQSEYETI KRYEKRKEQAANLNMTELVVKLEDMIADETNH MEELDRLLNDKAMVLN |
| 3IQ1 | *Vibrio cholerae* | Gram-negative, facultative anaerobic bacteria | 12 | Dps | MATNLIGLDTTQSQKLANALNNLLANYQVFYM NTRGYHWNIQGKEFFELHAKFEEIYTDLQLKI DELAERILTLSARPMHSFSGYLKAAQIKEHTDS IDGRSSMQGLVDGFSILLHQQRDILELAGETG DEGTSALMSDYIREQEKLVWMLNAWLK |
